# A universal, genome-wide guide finder for CRISPR/Cas9 targeting in microbial genomes

**DOI:** 10.1101/194241

**Authors:** Michelle Spoto, Elizabeth Fleming, Julia Oh

## Abstract

**Background:** The CRISPR/Cas system has significant potential to facilitate gene editing in a variety of bacterial species. CRISPR interference (CRISPRi) and CRISPR activation (CRISPRa) represent modifications of the CRISPR/Cas9 system utilizing a catalytically inactive Cas9 protein for transcription repression or activation, respectively. While CRISPRi and CRISPRa have tremendous potential to systematically investigate gene function in bacteria, no pan-bacterial, genome-wide tools exist for guide discovery. We have created Guide Finder: a customizable, user-friendly program that can design guides for any annotated bacterial genome.

**Results:** Guide Finder designs guides from NGG PAM sites for any number of genes using an annotated genome and fasta file input by the user. Guides are filtered according to user-defined design parameters and removed if they contain any off-target matches. Iteration with lowered parameter thresholds allows the program to design guides for genes that did not produce guides with the more stringent parameters, a feature unique to Guide Finder. Guide Finder has been tested on a variety of diverse bacterial genomes, on average finding guides for 95% of genes. Moreover, guides designed by the program are functionally useful—focusing on CRISPRi as a potential application—as demonstrated by essential gene knockdown in two staphylococcal species.

**Conclusions:** Through the large-scale generation of guides, this open-access software will improve accessibility to CRISPR/Cas studies for a variety of bacterial species.

## Background

The CRISPR/Cas system represents a considerable development in gene editing technology for a wide variety of organisms. Sequence-specific targeting is possible through interactions between a complementary guide RNA and the target sequence, and between the protospacer adjacent motif (PAM) and the Cas nuclease. At the target sequence, the Cas nuclease induces a double stranded break which is subsequently repaired by the cell using non-homologous end-joining (NHEJ) if it exists. This often results in deleterious insertion or deletion mutations that can disrupt the function of the target gene.

Given Cas9’s broad activity and efficacy, the CRISPR/Cas system been used to successfully edit genes across a diverse range of species[1][2][3], but its application to bacterial genome editing has been more limited. For instance, many bacterial species do not possess the machinery to efficiently repair double stranded breaks, and targeting with CRISPR/Cas is consequently lethal to the cell. Additionally, homologous recombination (HR)-mediated repair requires introduction of a second template either as linear DNA or on a supplemental plasmid. Nevertheless, the CRISPR/Cas9 system has significant potential to facilitate gene-targeting/editing in wide range of microorganisms[4]. Moreover, additional tools that do not depend on HR or NHEJ for disrupting gene function have since been developed, including CRISPR interference (CRISPRi) and CRISPR activation (CRISPRa).

CRISPRi and CRISPRa are modifications of the CRISPR/Cas system that employ a catalytically inactive Cas9 protein (dCas9) for targeting[5]. In the case of CRISPRi, the dCas9 is used for transcriptional repression by sterically blocking transcription machinery and preventing initiation or elongation, depending on the location of the target sequence on the promoter or DNA strand. CRISPRa works similarly, except it is fused to the omega subunit of RNA polymerase, allowing increased recruitment of the polymerase when targeted to sequences upstream of the -35 box of the promoter[5].

For all systems, the efficiency of targeting as well as the occurrence of off-target effects elsewhere in the genome is influenced by guide selection. A guide’s distance from transcription start site[6], GC content[7], homopolymer content[8], and cross-reactivity to similar sequences in the genome have been shown to affect targeting efficacy[6]. While these guide design constraints are important for efficient targeting, the consideration of these multiple factors during guide selection makes manual guide design impractical in large scale. This is of particular importance, for example, in genome-wide CRISPRi and CRISPRa studies, which require the design of thousands of guides[9].

Existing tools for guide design are limited in their generalizability to large numbers of diverse microbial genomes, which can vary widely in GC content, length, and number of repeat regions. Indeed, the majority of sgRNA design tools have been developed exclusively for eukaryotes or a small handful of model organisms[10][11][12][13][14]. Other programs possess flexibility for the input genome but are limited by a lower-throughput design[15], absence of user-defined filtering parameters, inability to automatically iterate with relaxed design parameters[15][16], or lack a user-friendly design[16]. Finally, as the efficacy of CRISPRi and CRISPRa are likely augmented by targeting multiple loci simultaneously within the same gene, identifying and optimizing multi-guide design is needed. No open-source guide designer combines the customizability and flexibility of user-defined design constraints, pan-bacterial applicability, gene iteration, and paired guide selection in a user-friendly format.

Thus, we have created Guide Finder to address these limitations. Our program has a simple input for any annotated complete or draft genome and accepts default or user-defined guide design parameters, which is important given the broad characteristics of different microbial genomes like GC content, size, or the presence of repetitive regions. Finally, the automated and iterative guide design is capable of designing guides to target any number of genes for any annotated bacterial genome, including optimizing selection of multiple guides for double targeting. Focusing on its applications for CRISPRi, we have demonstrated its utility in selecting guides genome-wide for a diverse set of bacterial species and its ability to select functional guides suitable for gene knockdown. Guide Finder is the first publically available, automated guide selection program designed specifically for bacteria that incorporates user-defined filtering parameters, off-target searching, and iterative guide design with utility for both complete and draft genome annotations. This tool will help facilitate flexible, large-scale guide design and thus improve access to high-throughput studies of gene function.

## Implementation

Guide Finder is written in the R programming language and is available free to use. Guide Finder was written such that it can be used to find guides for both complete and draft genomes, recognizing that many users may not have a complete genome for their organism of interest. The workflow of the program, including inputs and outputs, is described (Fig. 1).

**Figure 1.**
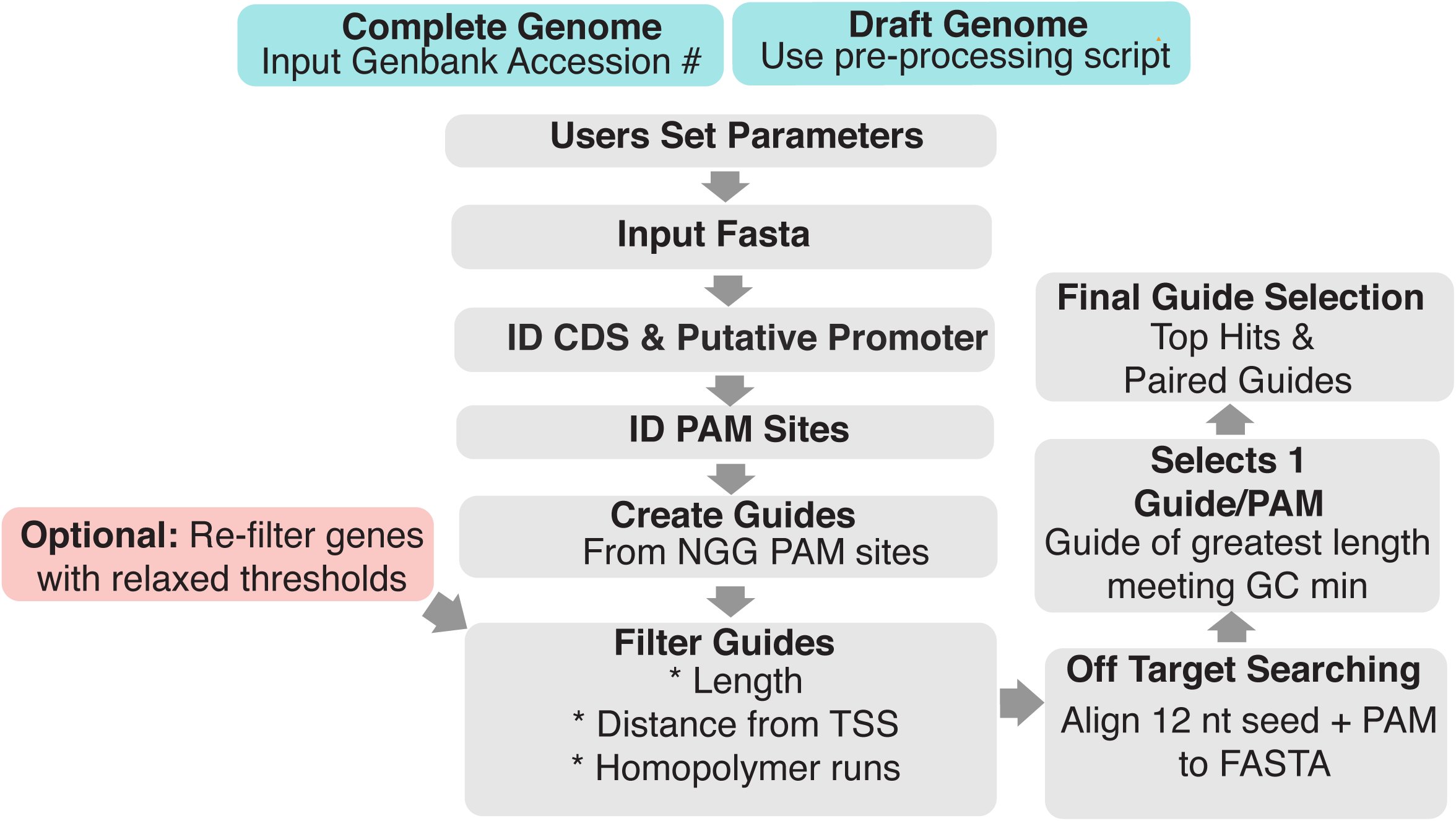
Guide Finder Workflow. Users set parameters and input FASTA file. Coding sequence and promoter coordinates are identified and used to obtain sequences. PAM sites are identified guides created and filtered. Off-target searching is conducted using BLAST. Final guide selection creates a Top Hits list and Paired Guides list. Genes without guides are identified and re-run with relaxed parameters. (PAM = protospacer adjacent motif, TSS = transcription start site, nt = nucleotide)

### Inputs & Outputs

#### Inputs

Guide Finder is capable of designing guides for both complete and draft genomes, although the inputs differ slightly.

#### Complete Genome

For complete genomes, users simply supply the Genbank accession number and fasta file.

#### Draft Genome

Given the variable organization and notation of draft genomes, annotated draft genome files must be preprocessed prior to input into the program. Utilizing the supplied pre-processing script, multi-sequence fasta files (e.g. fasta files containing sequence information for multiple contigs) must be concatenated into a single sequence, with the addition of a series of N’s between contigs. The coordinates of the coding sequences are then identified by aligning the coding sequences against the concatenated fasta file using BLAST and adjusted to the format required by the main Guide Finder script(i.e. the smaller coordinate designated as the “start” coordinate). These coordinates are then input into the main script, along with the single-sequence fasta file.

### Outputs

There are two main outputs of the guide finder program: Top Hits and Paired Guides lists. Intermediate outputs, such as a list of all possible unfiltered guides, are also made available to the user for reference.

#### Top Hits List

A list of guides preferentially selected based on their proximity to the transcription start site. The maximum number of guides supplied per gene is set by the user.

#### Paired Guides List

A list of guide pairs, designed to doubly target the same gene in the same cell to increase targeting efficiency. Suitable guide pairs are selected on the basis of the distance between the guides, a parameter set by the user.

### Program Workflow

#### Coordinate Identification

The identification of gene start and end coordinates is the first step in the Guide Finder workflow and differs slightly for complete versus draft genomes. For complete genomes, the script reads in the annotated genome file containing the gene coordinates and modifies the coordinates to include the putative promoter region. For draft genomes, the coordinates—identified during pre-processing—are directly input into the program and modified to include the putative promoter region.

#### Coding and Promoter Sequence Retrieval

The gene start and end coordinates are used to retrieve the coding and putative promoter sequences from the fasta file.

#### Guide Creation

Searching within the promoter and gene body, the program identifies NGG PAM sites and utilizes the sequence around each site to create three guides/PAM site (of length 20 bp, 21 bp, and 22 bp.) The varied guide length selection increases the number of potential guides, many of which will be lost to filtering, as described below.

#### Guide Filtering

Guides are filtered according to default and user-defined parameters. By default, the program removes any guides that contain a homopolymer run of As or Ts and guides of inadequate length (<20 bp). A user-set threshold is used to filter based on maximum distance from the start site, as targets closest to the transcriptional start site are most likely to disrupt gene function. Guides are also filtered to minimize potential off target effects. The first 12 nucleotides closest to and including the PAM site for each guide is aligned to the fasta file and guides that match to more than one location in the genome are discarded.

#### Final Guide Selection

For each PAM site, the program selects the guide of the greatest length that meets GC minimum set by the user. From these guides, two final guides lists are created: Top Hits and Paired Guides, which provide guides and guide pairs suitable for single and dual gene knockdown, respectively.

#### Iteration

The program identifies genes that did not produce any guides with the primary parameters. Users have the option to lower these thresholds and re-run these genes through the program to identify additional guides. Users can elect to reduce the GC minimum, increase the maximum guide distance from the transcription start site, retain guides that contain homopolymers, and relax off target searching. Users can relax each of these guide design constrains individually or in combination.

### Results & Discussion

Guide Finder is intended to reduce the effort required to design guides targeting genes in any bacterial species and accommodates both complete and draft genome annotations, the latter of which is important given the large number of unique isolates being sequenced and investigated. The program is customizable and incorporates user-defined guide constraints, including: minimum GC content, maximum distance from the transcription start site, and distance between guides (for dual targeting knockdown).Recognizing the diversity of bacterial species, we aimed to create a program where users could tailor guide design parameters based on the characteristics of their organism of interest, for example, setting a relatively low guide GC minimum while working with a GC poor species. Additionally, the program identifies genes for which no guides meeting set thresholds could be identified, allowing iterative guide-calling to maximize the number of genes targeted. Users have the option to re-run these genes through the guide finder program with relaxed design constraints to identify additional guides.

Although users can elect to design guides for just one gene or a handful of genes, if desired, the program is intended to be particularly useful for large-scale guide design. To investigate these intended uses, we conducted tests *in silico* and *in vitro* focusing on CRISPRi to determine: 1) the utility of the program across diverse bacterial species and 2) the ability of the program to design functional guides.

### Guides for diverse genomes

#### Testing on Complete Genomes

Guide Finder was utilized to create guides across the genome for a diverse set of ten complete bacterial genomes (Table 1). These genomes were selected for their diversity in genome size, percentage of gene duplications, and GC content. For each genome, preliminary parameters were set as: a GC minimum of 35%, a maximum distance from the TSS of 30%, and a minimum distance between guides of 100 base pairs, based on the projected footprint of the Cas9 protein[17]. These parameters have been utilized in our lab previously for successful gene knockdown with CRISPRi and thus represent rational design constraints. For each genome, genes that did not produce suitable guide pairs or single guides were identified by the program. These genes were re-run with the following constraints: a GC minimum of 30%, a maximum distance from the TSS of 50%, retention of guides with homopolymers, and relaxed off-target searching. These parameters were relaxed individually and in combination. The guide finder program was able to successfully select guides for each of the diverse genomes irrespective of genome size or GC content, but differences in output and run-time were observed (Fig 2).

**Table 1.** Complete genomes tested. Ten complete genomes, obtained from NCBI, were selected for their varying genome size and GC content, as noted in this table.

| Phylum | Organism | Genome Size (Mb) | GC Content (percentage) |
| --- | --- | --- | --- |
| Firmicutes | *Lactobacillus brevis* | 2.29 | 45 |
|  | *Lactobacillus jensenii* | 1.67 | 34 |
|  | *Staphylococcus aureus* | 2.82 | 33 |
|  | *Staphylococcus epidermidis* | 2.49 | 32 |
| Proteobacterium | *Acinetobacter baumannii* | 4.33 | 39 |
|  | *Rhizobium leguminosarum* | 4.85 | 60 |
|  | *Pseudomonas aeruginosa* | 6.26 | 66 |
| Actinobacterium | *Mycobacterium tuberculosis* | 4.41 | 66 |
|  | *Micrococcus luteus* | 2.50 | 73 |
|  | *Streptomyces scabiei* | 10.41 | 71 |

**Figure 2.**
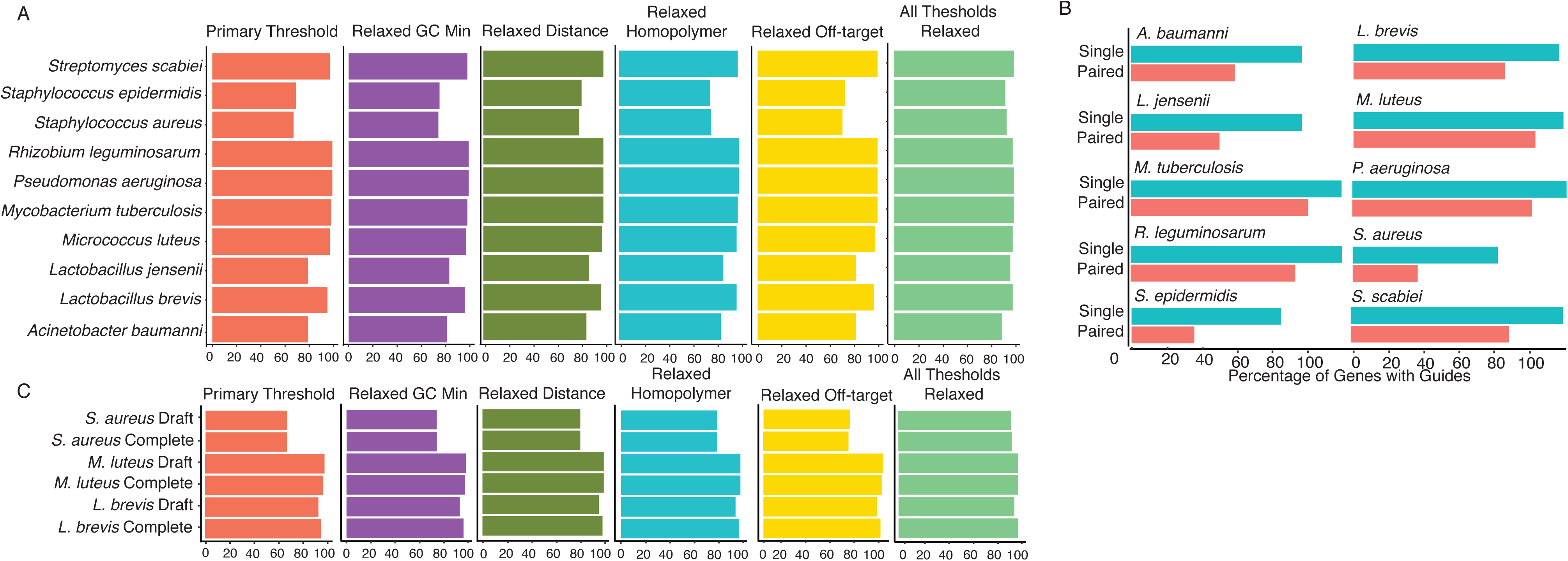
Testing on Complete and Draft Genomes *in silico.* **A. Complete Genomes.** Guide Finder was tested on 10 complete genomes under primary design constraints with iteration under relaxed constraints (individually and in combination). **B. Complete vs Draft.** Guide Finder was tested on 3 draft genomes; percentage of genes with guides was compared to a complete genome of the same species. **C. Paired vs Single Guides.** The percentage of genes targeted by single versus paired guides was compared.

### GC Content

As expected, genomes with lower GC content (<40%) were less successful in producing usable guides for each gene. For *S. epidermidis, S. aureus*, *A. baumanni*, and *L. jensenii* genomes (GC contents of 33%, 32%, 39%, and 34%, respectively), the percentage of genes producing guides under the primary filtering thresholds was considerably lower than the average for all ten genomes (87.5%) at 68%, 67%, 79%, and 79%, respectively. The average for genomes > 40% GC content was 97.5%. However, for genomes with low GC content, iteration with lowered parameters was very useful in recovering genes that did not originally produce guides. When each design constraint was relaxed in combination, the percentage of genes with guides improved to 98%, 93%, 89%, and 96% for *S. epidermidis, S. aureus*, *A baumanni*, and *L. jensenii*, respectively (Fig. 2A).

#### Gene Duplications

We hypothesized that a genome known to contain a high percentage of gene duplications, such as *Mycobacterium tuberculosis,* would have difficulty producing a large number of usable guides, owing to the high probability of off-target matching. Surprisingly, however, this genome was able to create guides for 98% of genes using primary thresholds, probably owing to its relatively high GC content (65%).

#### Genome Size

Although Guide Finder was run successfully on each of the ten genomes tested, runtime increases with genome size due to the increased number of genes and subsequent increased number of potential guides (each of which is analyzed for GC content, location, etc.). For example, the program takes approximately 10 minutes to complete using the *S. epidermidis* genome (2.49 Mb) but takes approximately 18 hours for the largest genome tested, *S. scabeii* (10.41 Mb). *S. scabeii* is one of the largest known bacterial genomes and thus we do not expect that this issue will affect most users but represents a potential area of improvement for future versions of Guide Finder.

#### Double Guide Design

Guide Finder is capable of designing guides for multi-guide targeting, which may improve efficacy of knockdown. Aside from the fact that overlapping guides have been shown to reduce knockdown efficiency, very little is known about the impact of the distance between dual targeting guides on gene knockdown in bacteria[6]. However, it is plausible that the footprint of the Cas9 protein may influence the ability of two nearby guides to target simultaneously. For this reason and to allow flexibility as new information becomes available, Guide Finder allows users to set a minimum distance threshold that guides selected for dual knockdown must meet. As expected, paired guide creation—including a 100 bp distance-between-guides threshold—is feasible for fewer genes than single guide creation, owing to the fact that some genes may produce only a single suitable guide or produce guides that are located in close proximity (Fig.2B).

### Testing on Draft Genomes

Three draft genomes were selected to test utility for incomplete genome annotations and compared to a complete genome annotation of the same species. Draft annotations were obtained from the Pathosystems Resource Integration Center (PATRIC)[18] and whole-genome nucleotide sequences and coding sequences for incomplete genomes were obtained from NCBI. Incomplete genomes were pre-processed with the supplied script to identify gene coordinates. Incomplete genome annotations were successfully used to design guides across the genome for each of the three species tested. In terms of percentage of genes with identified guides and run-time, there are no appreciable differences between complete and incomplete genome annotations (Fig. 2C). This result highlights the utility of the program for both types of genome annotation files.

### Essential gene knockdown to validate guides

We evaluated the functional utility of Guide Finder guides by random assessment of essential gene knockdown in *Staphylococcus (S.) aureus* and *S. epidermidis,* focusing on CRISPRi as a potential application. Nearly all guides showed effective knockdown manifested as growth defects with the exception of *groEL* and *rpoC* (Fig. 3). Further investigation measuring transcription of the locus using qPCR showed that the guide targeting *rpoC* did not reduce transcription (highlighting the value of predicting and testing multiple guides). *groEL* was effectively targeted but was either non-essential under our tested condition, or residual transcript could be rescuing cell function (Fig. 4). Thus, our Guide Finder parameters have been used for successful gene knockdown and thus represent rational design constraints. Overall, these results highlight the utility of the guide finder program to create functional guides—demonstrated by functional testing of essential genes—and underscores the need for continued investigation of guide design for improved targeting efficacy in bacterial species. With customizable, user-defined design parameters and access to program source code, users are able to adjust guide selection as this information becomes available.

**Figure 3.**
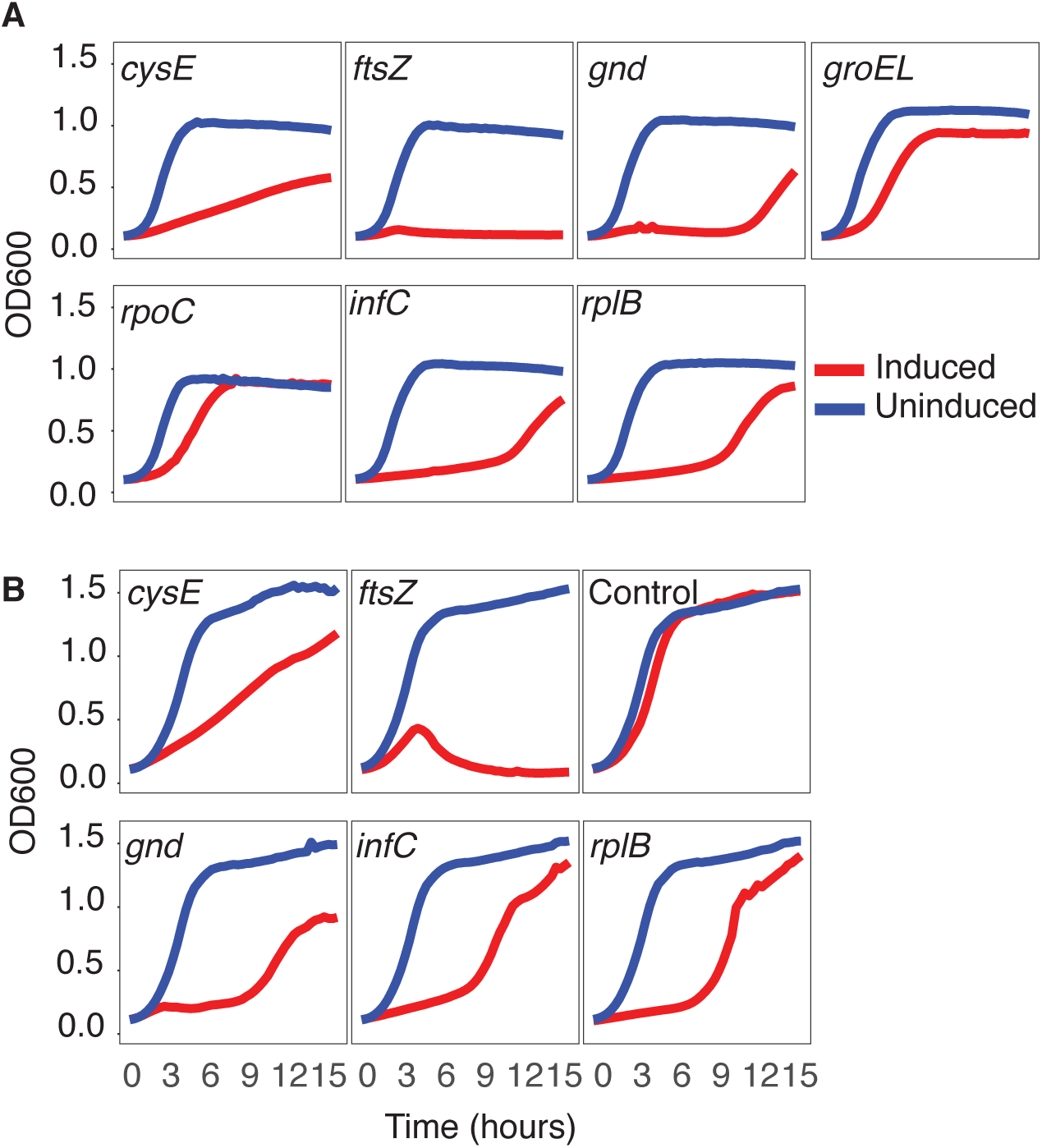
Essential gene knockdown. Essential genes were targeted for knockdown in *S. aureus* (A) and *S. epidermidis* (B) and growth curves were created from OD measurements over a 16 hour growth assay. ATc= anhydrotetracycline induction, uninduced = control. Control: empty vector (no guide) acts as a control, indicating that the growth defect is not due to ATc administration. With the exception of *groEL* and *rpoC*, the knockdown of most essential genes caused a growth defect, as expected.

**Figure 4.**
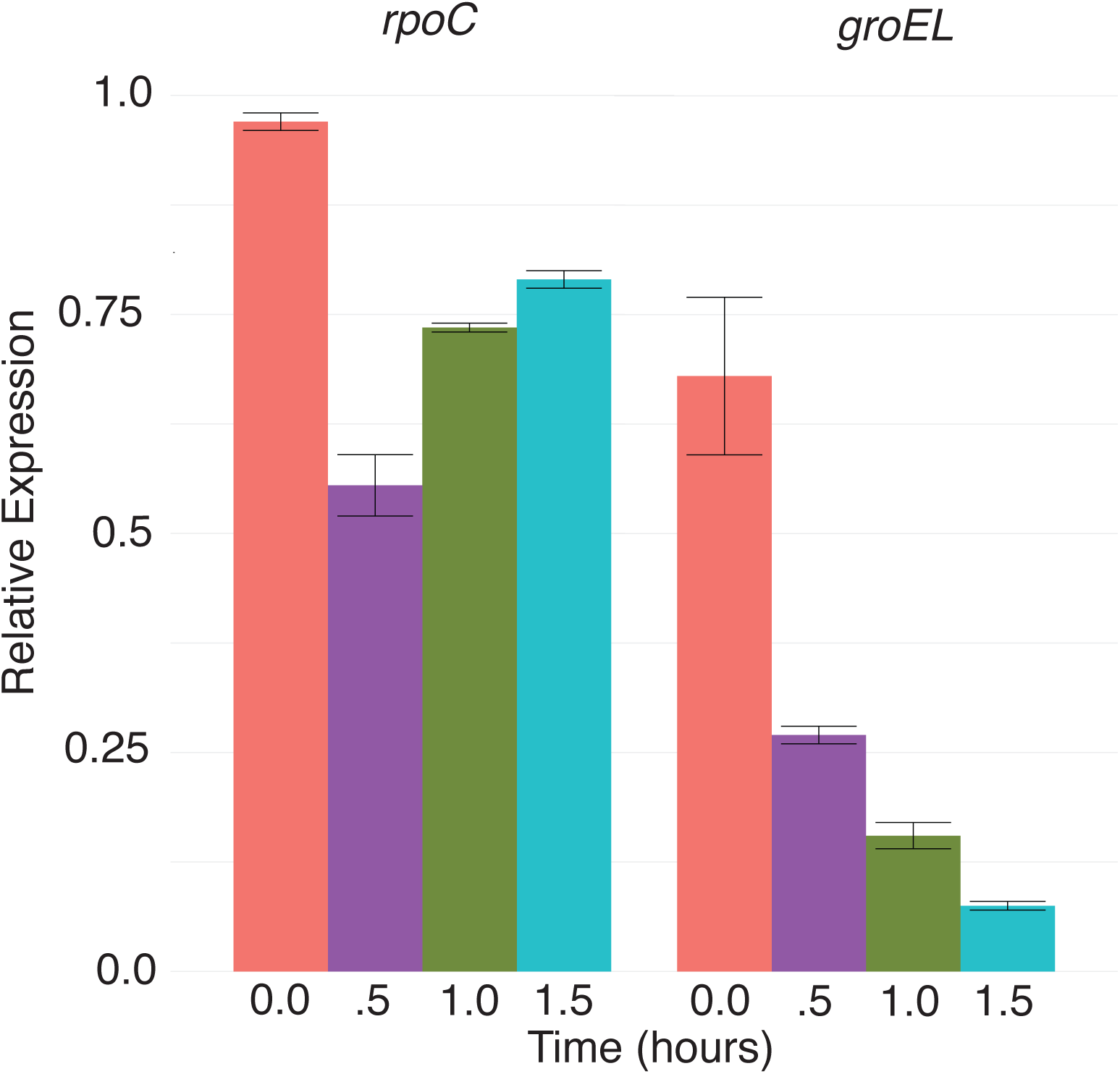
*rpoC* and *groEL* Relative Expression. mRNA levels of *rpoC*and *groEL* were measured in knockdown strains over a 1.5 hour growth assay. Transcript levels were normalized to the control strain, at each time point.

## Conclusions

As the first user-friendly pan-bacterial automated program suitable for large-scale guide selection, this guide finder program is capable of designing guides for any number of genes for any annotated bacterial genome. Guide Finder provides users with a ready-to-use list of designed guides without the need for gene-by-gene score comparison or additional filtering. In this way, Guide Finder’s utility lies in its ability to not only design guides in a large-scale format but to also provide users with the most-suitable guides for each input gene, according to the parameters they defined. By enabling high quality, large-scale guide selection for any bacterial genome, Guide Finder improves access to high-throughput studies of bacterial gene function, including genome-wide CRISPRi and CRISPRa studies.

## Availability and requirements

**Project name:** Guide Finder

**Project home page:** https://github.com/ohlab/Guide-Finder

**Operating system(s):** Mac, Windows

**Programming language:** R

**Other requirements:**

**License:** None

**Any restrictions to use by non-academics:** None

## Abbreviations

CRISPRi: CRISPR interference
CRISPRa: CRISPR activation
PAM: protospacer adjacent motif
NHEJ: non-homologous end-joining
HR: homologous recombination
ATc: anhydrotetracycline
Cm: chloramphenicol
TSB: tryptic soy broth

## Declarations

**Ethics approval and consent to participate**

Not Applicable

## Consent for publication

Not Applicable

## Availability of data and material

Data sharing is not applicable to this article as no datasets were generated or analyzed during the current study. Genomes used for analysis were obtained from NCBI and PATRIC. Specific strains (including accession and genome ID numbers) are listed in the supplemental material.

## Authors' contributions

JO identified the need for a large-scale guide finder program in bacteria and provided feedback and guidance during the development of the program. MS developed the pre-processing and guide finder scripts, performed all *in silico* experiments and analyses, assisted EF in performing the *in vitro* studies and analyses, and wrote the manuscript.EF performed the *in vitro* studies and analyses. All authors read and approved the final version of this manuscript.

## Acknowledgements

The authors thank The Jackson Laboratory for Genomic Medicine Computational Sciences department for their assistance in code optimization and manuscript review.

## Supplemental Information

Methods for strain knockdown creation, growth assays, and transcript measurements are detailed below.

## Knockdown Strain Creation

For both species, knockdown strains were created as follows: For each targeted gene, a single guide was designed by the guide finder program for targeting. The guide was ligated into our custom CRISPR/dCas9 shuttle vector. Our CRISPR/dCas9 shuttle vector includes all of the necessary components for CRISPRi, including: dCas9 under an anhydrotetracycline (ATc) inducible promoter, dCas9 handle (crRNA and tracrRNA fusion), and a chloramphenicol resistance maker (for selection). The shuttle vectors containing the proper targeting guides were transformed into *E. coli* and resultant colonies screened for the guide sequence. A single positive colony was grown in TSB with chloramphenicol (TSM/Cm) overnight and, using the QIAprep Spin Miniprep kit,plasmids were isolated from *E. coli* and transformed into our Staphylococcal species of interest. For *S. aureus,* plasmids were transformed via electroporation into competent *S. aureus* RN4220 cells. For *S. epidermidis,* phagemid transfer was utilized to incorporate the plasmid into *S. epidermidis* strain Tu3298, according to the protocol described elsewhere[19].

## Growth Assays

Growth assays were performed to assess knockdown of essential genes. Growth assays were performed in both *Staphylococcus aureus* and *Staphylococcus epidermidis,* as follows: A single colony of each knockdown strain was grown in TSB containing chloramphenicol overnight. The overnight culture was diluted to an OD of 0.05 in TSB/Cm, grown to an OD of 0.5, and diluted again at the start of the assay to an OD of 0.05 with TSB/Cm (control group) or TSB/Cm + 0.1 uM anhydrotetracycline (inducer, experimental group). The cultures were grown for 16 hours, with OD measurements taken each half an hour to construct a growth curve for each knockdown strain. For each strain, the induced/experimental group growth curve was compared to the uninduced/control group curve. Knockdown of most essential genes resulted in a severe growth defect, as expected. The knockdown of two genes, *groEL* and *rpoc,* did not result in the expected growth defect and we investigated the ability of each guide to reduce transcript levels.

## Measuring Transcript Levels

In *S. aureus,* we measured transcript levels of *groEL* and *rpoc* growing in liquid media to determine if the selected guide was capable of reducing transcript levels. A single colony of each *groEL* and *rpoc* knockdown *S. aureus* strain was grown overnight in TSB/Cm at 37 C, shaking. The overnight culture was back diluted to an OD of 0.05 and grew up at 37 C until an OD600 of 0.5. The culture was back diluted again to an OD of 0.05 with TSB containing chloramphenicol and 0.1uM anhyotetracycline and were grown for 1.5 hours; time points were taken throughput the assay at hours 0, 0.5, 1, and 1.5. An aliquot taken at each time point was mixed with 2 volumes of RNA protect and incubated for 5 minutes at room temperature. The aliquot was spun down, supernatant decanted, and stored at -20 until RNA extraction. RNA from the four time points was extracted according to the protocol for the RNaeasy Plus kit with an added enzymatic digestion using lysozyme and lysostaphin, for lysis of the Gram positive *S. aureus*. RNA was reversed transcribed to create cDNA using the High-Capacity cDNA Reverse Transcription kit (Applied Biosystems), according to provided instructions. QPCR was performed using PowerUp™ SYBR^®^ Green Master Mix (Applied Biosystems) in conjunction with gene specific primers. Primers amplifying the gene ftsZ were used as an internal control and non-template controls were included. Duplicate QPCR reactions were performed for each assay as a technical replicate.

## Genomes Used in Analysis

Genomes used for complete genome analysis were obtained from NCBI. Accession numbers for each strain is listed below:

*Lactobacillus brevis:* CP000416.1

*Lactobacillus jensenii:* CP018809.1

*Staphylococcus epidermidis*: AE015929.1

*Staphylococcus aureus*: CP000253.1

*Rhizobium leguminosarum*: CP007045.1

*Pseudomonas aeruginosa*: AE004091.2

*Mycobacterium tuberculosis*: AL123456.3

*Micrococcus luteus*: CP001628.1

*Streptomyces scabiei*: FN554889.1

Genomes used for draft genome analysis were obtained from PATRIC. Strain used and genome ID number is listed below.

*Micrococcus luteus ATCC 12698*. **Genome ID:** 1270.61

*Micrococcus luteus O’kane.* **Genome ID:** 1270.50

*Staphylococcus aureus WBG10049*. **Genome ID:** 585160.3

*Staphylococcus aureus SA14-296.* **Genome ID:** 46170.233

*Staphylococcus epidermidis NLAE-zl-G239*. **Genome ID:** 1282.2004

*Staphylococcus epidermidis FDAARGOS_83*. **Genome ID:** 1282.1163

